# Improving intermolecular contact prediction through protein-protein interaction prediction using coevolutionary analysis with expectation-maximization

**DOI:** 10.1101/254789

**Authors:** Miguel Correa Marrero, Richard G.H. Immink, Dick de Ridder, Aalt D.J van Dijk

**Affiliations:** Bioinformatics Group, Wageningen University, Wageningen, The Netherlands; Laboratory of Molecular Biology, Wageningen University & Research, Wageningen, The Netherlands; Bioscience, Wageningen University & Research, Wageningen, The Netherlands; Biometris, Wageningen University, Wageningen, The Netherlands

## Abstract

Predicting residue-residue contacts between interacting proteins is an important problem in bioinformatics. The growing wealth of sequence data can be used to infer these contacts through correlated mutation analysis on multiple sequence alignments of interacting homologs of the proteins of interest. This requires correct identification of pairs of interacting proteins for many species, in order to avoid introducing noise (i.e. non-interacting sequences) in the analysis that will decrease predictive performance. We have designed Ouroboros, a novel algorithm to reduce such noise in intermolecular contact prediction. Our method iterates between weighting proteins according to how likely they are to interact based on the correlated mutations signal, and predicting correlated mutations based on the weighted sequence alignment. We show that this approach accurately discriminates between protein interaction versus noninteraction and simultaneously improves the prediction of intermolecular contact residues compared to a naive application of correlated mutation analysis. Furthermore, the method relaxes the assumption of one-to-one interaction of previous approaches, allowing for the study of many-to-many interactions. Source code and test data are available at www.bif.wur.nl/

## Introduction

Virtually any biological process requires, at some point, recognition and interaction between specific proteins. Knowing which residues mediate a protein-protein interaction (PPI) can lead to better understanding of its interaction specificity. It is also useful for modeling protein complex structures (1) and for designing drugs targeting PPIs (2). Unfortunately, experimentally elucidating which residues make contact across interfaces (by, for example, alanine scanning or cocrystallization of the proteins of interest) is low-throughput and laborious. Hence, a plethora of computational methods have been designed for this task. Amongst these, coevolutionary analysis has received much attention in recent years, thanks to the expanding wealth of sequence data and methodological advances.

Coevolutionary methods, applied to the study of interacting proteins, use sequence information alone to predict contacts between proteins. They rely on the emergence of shared evolutionary constraints between interacting proteins to maintain the interaction, which can be observed as correlated mutations (3). Revealing these correlated mutations requires building large multiple sequence alignments (MSAs) of interacting homologs of the proteins under study, which are then analyzed using a variety of methods (Figure 1a). These two MSAs are paired: each row needs to contain a pair of interacting proteins. However, it is not trivial to avoid introducing non-interacting pairs of sequences, which adds noise to the analysis and decreases predictive performance (Figure 1b), a widely recognized problem in the field (4–6). This can be caused, for example, by gene duplication events, which might result in divergence of interaction patterns in the resulting proteins (7, 8). When analyzing prokaryotic proteins, this is usually dealt with by filtering by genomic colocalization: a protein will often interact with a paralog within the same operon, and not with those outside. This does not hold true for eukaryotic proteins, where coevolutionary analysis has been a challenge.

**Fig. 1.**
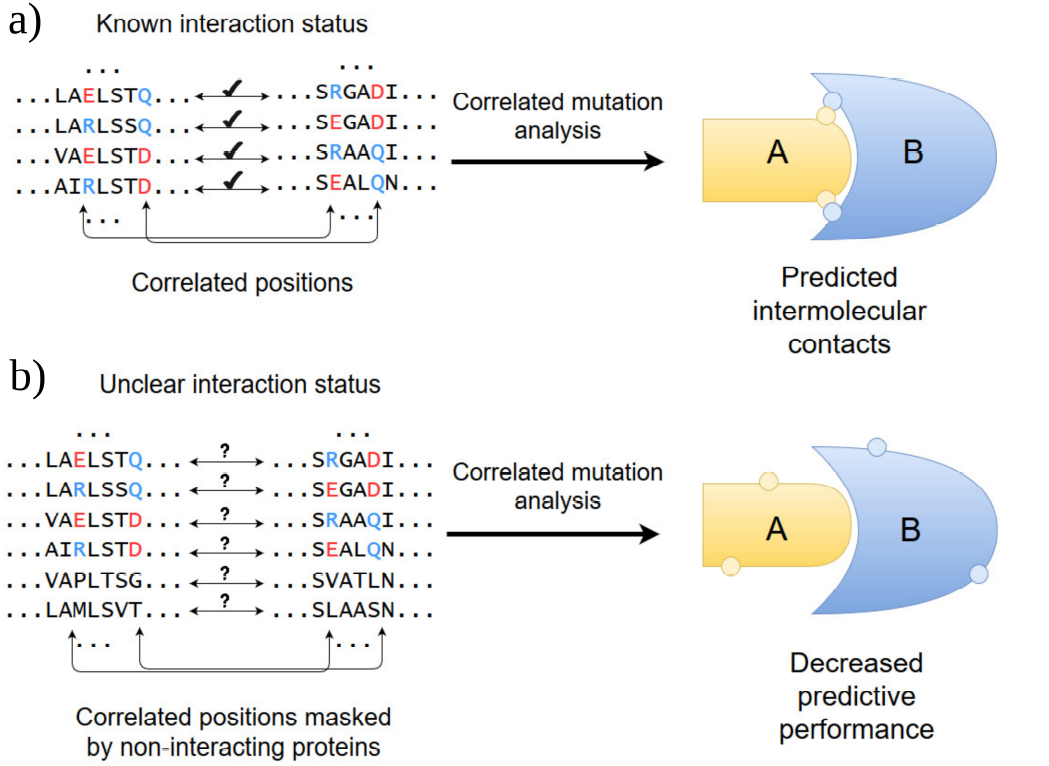
Contact prediction between hypothetical proteins A and B by coevolutionary analysis of their paired MSAs. (a) Ideal case, where sequences in the MSAs can be matched by their interaction status. This is approximated when analyzing prokaryotic proteins by matching them by genomic colocalization. Circles indicate residues predicted to participate in the interaction. (b) Case where interaction status is unknown. Inadvertent introduction of non-interacting sequences (two last pairs of sequences) leads to decreased contact prediction performance.

Before the appearance of coevolutionary methods which remove indirect relationships between residues (9), approaches to predict PPIs based on MSAs using coevolutionary information had already been developed (10, 11), although they were not applied to improve contact prediction. Algorithms that address both removal of indirect relationships and PPI prediction have only appeared recently (12, 13). These methods aim to maximize the coevolutionary signal by simultaneously matching pairs of interacting proteins within a species and predicting the intermolecular contacts, with no *a priori* knowledge of either.

In this paper, we present Ouroboros, a new approach to concurrently predict protein-protein interactions and intermolecular contacts, based on the expectation-maximization (EM) algorithm. It relies on iteratively improving two models, one of protein coevolution and another of independent evolution, in order to discriminate between interacting and noninteracting proteins and thus improve the coevolutionary signal. As the previously mentioned methods, it does so with no prior information about protein interactions or intermolecular contacts. Algorithmic and statistical considerations aside, our approach differs from the work of Bitbol et al. and Gueudré et al. in that it relaxes their assumptions. Most importantly, both papers consider that a protein can only interact with one protein (one-to-one interactions). Ouroboros, as we show here, allows for many-to-many interactions. This is particularly relevant given the prevalence of many-to-many interactions in eukaryotes, for example, between members of large protein families (Van Wijk et al. 14, Immink et al. 15, Reinke et al. 16).

We apply the newly developed algorithm on a dataset containing many-to-many interactions and non-interacting proteins. We show that Ouroboros is able to accurately discriminate between interacting and non-interacting proteins using only coevolutionary information. In turn, this allows it to improve intermolecular contact predictions compared to a naive application of coevolutionary analysis.

## Materials and Methods

### A. Data representation

Consider two multiple sequence alignments (MSAs) *M*_1_ and *M*_2_, both containing *S* sequences of length *L*_1_ and *L*_2_, respectively. Each row *i* of one of the matrices represents an aligned protein sequence. The row *i* in the other matrix represents a protein sequence that might interact with it. We will call these sequence pairs. We transform the matrices *M*_1_ and *M*_2_ into the matrices *N*_1_ and *N*_2_, where each element of the original matrices is mapped to numbers in the set *A* ∊ {0,…,20}, which represent the 20 standard amino acids plus the gap symbol (-). We will refer to matrices *N*_1_ and *N*_2_ as numerical matrices, and their columns serve as response vectors in our modeling approach.

The matrices *N*_1_ and *N*_2_ are, in turn, expanded to matrices *B*_1_ and *B*_2_, containing *S* rows and *L*_1_ · (|*A*| – 1) resp. *L*_2_ · (|*A*| – 1) columns. Each element of a numerical matrix is turned into a set of |*A* –| elements of a binary matrix, where a 1 indicates the amino acid present at a certain position; gaps are denoted by zeros. We shall refer to matrices *B*_1_ and *B*_2_ as binary matrices. These are used as explanatory variables in our coevolutionary model.

Columns with a gap frequency equal to or greater than 0.5 are excluded from the analysis, as they yield insufficient information. Constant columns are also removed, given that the method relies on finding covariant signals.

### B. Modeling setup

Each sequence pair *i* in the alignments is associated with a hidden variable *z_i_*, whose value indicates a probability of interaction. To discriminate between interaction and non-interaction, the algorithm uses two different models, which are described below. These models are weighted by the values of *z*. To assign initial values to *z*, we give each sequence pair the maximum weight possible in both models described below, and simply follow the procedure described in this section to estimate the model parameters and assign values to *z*. Subsequently, the EM algorithm is applied, which iteratively applies two steps until convergence: the two models are refined in the M step and new values of *z* are derived in the E step.

#### B.1. Coevolutionary model

We break down searching for covariation between the two MSAs into a number of multiclass classification problems. We model each column of each MSA as a function of the sequences in the other MSA using multinomial logistic regression, where each amino acid constitutes a separate class. For each column in an MSA, the response variable is the corresponding column of the pertinent numerical matrix (*N*_1_ or *N*_2_); the explanatory variables are contained in the binary matrix that represents the other protein (*B*_2_ or *B*_1_). Each sample is weighted by the associated hidden variable value *Z_i_*. As we are only interested in intermolecular contacts, intramolecular covariation is not taken into account in our model. In order to prevent overfitting, the models are regularized using the elastic net penalty, which linearly combines L1 and L2 penalties. Based on previous work, the value of the mixing parameter is set to 0.99 (17). This value emphasizes the L1 penalty. In our implementation, models are fitted and regularized using the SGDClassifier in scikit-learn v0.19 (18).

The elastic net has an *α* parameter which controls the strength of the regularization, which must be tuned. This parameter is set independently for each column of the MSAs during model initialization. We train models for each column over a range of 15 *α* values spaced evenly on a log scale between 1 × 10^−3^ plus some weaker regularization strengths (from 10 to 40) for some extreme cases. We select a suitable value using the Bayesian Information Criterion (BIC) (19), which is computed as:

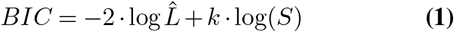

where log 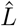 is the log-likelihood of the data according to the logistic model and *k* the number of model degrees of free-dom. The value of *α* that minimizes the BIC is chosen. Although the BIC penalizes model complexity, this penalty can easily be overcome by the goodness of fit achieved by simply adding a large number of explanatory variables that are unrelated to the learning task. Thus, using the BIC in a highdimensional setting can easily lead to overfitted models, an issue already described by several authors (see, for example, Chen and Chen 20 and Bogdan et al. 21). In order to avoid this, we set a threshold of 100 model degrees of freedom. This has a physical justification: a given residue will only interact with a limited number of residues. Once we have fitted logistic models for each column, we can calculate the posterior probability of each single residue according to them. Thus, we can compute the likelihood of a sequence pair *S_i_* under the coevolutionary model, which is given by

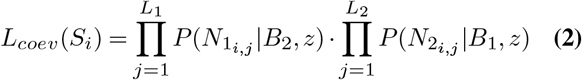

Here, the likelihood of a sequence pair is expressed as a product of two terms, one for each sequence of the pair. Each term is a product over the likelihoods assigned to each position (*N*_1_*i, j*__ or *N*_2_*i, j*__) according to the logistic regression models.

#### B.2. Model of independent evolution

The model of independent evolution considers that two given proteins evolve independently. Under this model, the probability of an amino acid is the frequency of the amino acid in its column, weighted by 1 – *z_i_*. Thus, the likelihood of sequence pair *S_i_* under the null model is given by

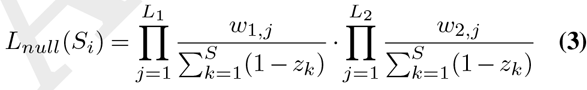

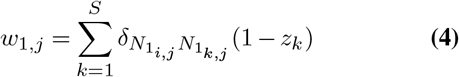

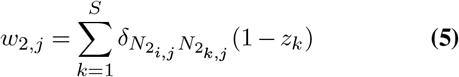

where *δ_ab_* denotes the Kronecker delta function, whose value is 1 if *a* = *b* and 0 otherwise.

##### B.3. Update of hidden variable values and convergence

In the E step, we compare the two models to estimate interaction probabilities. We derive new values of *z* using the equation

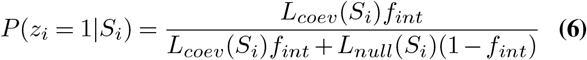

which can be derived from Bayes’ theorem. Here, *f_int_* is a prior fraction of interacting proteins. The procedure is repeated until the algorithm reaches convergence. We consider this happens when

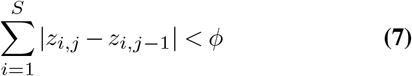

Here, *z_i, j_* indicates the current values of the hidden variables and *z_i, j−1_* those in the previous iteration, and *ϕ* is a userdefined convergence threshold (default 5 × 10^−3^).

#### C. Contact prediction

We have used the Julia implementation of plmDCA (Ekeberg et al. 22, 23) to predict intermolecular contacts on unfiltered alignments containing interacting and non-interacting sequence pairs (naive approach) and sequence alignments obtained from the EM-based approach described above.

#### D. Datasets

##### D.1. PDZ-peptide interaction dataset

We have used a dataset obtained by Tonikian et al. (24). In their study, the authors identified binding peptides for 82 different PDZ domains (54 human domains and 28 from C. *elegans*) from a C-terminal phage-displayed library of more than 10 billion random peptides. We have used a large subset of 3066 canonically binding peptides.

To evaluate PPI prediction performance, we need not only an interacting set, but also a non-interacting set. The dataset, however, contains no information about non-interaction. As done in previous work under the same circumstances (17, 25), we randomize the interacting set in order to create a noninteracting set. Nonetheless, randomly matching domains and peptides could generate a sizeable proportion of pairs that could actually interact. In order to minimize the chance of this happening, we incorporate prior knowledge regarding PDZ-peptide specificity detailed in the original paper, where the authors defined different specificity classes. Domains marked as having a unique specificity are grouped together with their closest neighbour. The exception is the PDLIM4-1 domain, which forms an outgroup in their classification scheme; we consider it as a separate class. To create the non-interacting set, domains are randomly selected and randomly matched with peptides different from those in their own specificity class. However, some classes share a few peptides, indicating these specificities have some small overlap. Thus, this procedure cannot guarantee that no interacting pairs are accidentally generated. As a result, a small fraction of pairs in the non-interacting set that we consider misclassified could actually interact.

To determine which pairs of residues are in contact between the PDZ domain and the binding peptide, we used four different complex structures previously used by the authors of the dataset (PDB IDs: 1BE9,1IHJ,1N7F,1N7T). We did not use the other structures they used, because the two sequences were fused and the observed contacts might be biologically irrelevant. We consider that two residues are in contact if there are 8Å or less between their Cβ atoms (or, if the residue is glycine, its Cα atom). To map these contacts to the sequences in the dataset, we align the sequences in the dataset and those of the structures using MAFFT v7.310 (26), which allows us to obtain a contact map between the domain and the peptide sequences. Contacts that appear in any of the structures are included in the contact map. The resulting alignments (without the sequences extracted from the PDB files) are used for coevolutionary analysis.

##### D.2. Synthetic data

We generated artificial MSAs containing both covarying sequence pairs (to simulate interaction) and non-covarying sequence pairs (to simulate non-interaction) in order to test the algorithm. The interacting pairs contain four columns that covary between the two sequences, where the residues in one column complement those in the other. On the contrary, non-interacting sequences contain residues drawn from the same distribution, but assigned at random, thereby removing the correlation between the sequences. Nonetheless, by chance, some non-interacting pairs may end up with a pattern of residues that resembles that of the interacting pairs. Non-interacting pairs that contain the pattern of residues that would be expected in an interacting protein in more than two columns are removed. Also, to simulate positions that do not participate in the interaction, these artificial MSAs contain columns with random distributions of amino acids. Each artificial sequence contains 15 positions in total.

## Results and discussion

### E. Combination of coevolutionary analysis with expectation-maximization

Naive application of coevolutionary analysis of MSAs containing both interacting and noninteracting proteins can lead to poor intermolecular contact prediction. The problem with such naive application is that we cannot properly estimate the parameters of the model describing residue couplings in a setting where the MSAs contain non-interacting proteins. On the other hand, if the parameters of the model were available, we should be able to identify pairs of sequences that do not follow these patterns of coevolution, which would indicate non-interaction.

A way to approach this kind of problem is the expectation-maximization (EM) algorithm. EM (27) is a general method for finding the maximum likelihood estimate of the parameters of a statistical model when the model depends on unobserved hidden variables (in this case, interaction or noninteraction). EM alternates between two steps: deriving the expected values for the hidden variables *z* based on the model parameters *θ* (E step), and re-estimating the model parameters *θ* based on the values of *z* (M step). In this work, we have combined a coevolutionary analysis algorithm (17) with EM. In this way, we can simultaneously model intermolecular contacts and interaction/non-interaction status. By weighting proteins in our models according to our predictions of their interaction status, we aim to boost the coevolutionary signal and improve predictions of intermolecular contacts.

To do so, we use two different models of protein evolution to distinguish interaction from non-interaction. The coevolutionary model considers that there is covariation between the two proteins, which points to interaction. The null model, however, assumes that the two proteins evolved independently, which would indicate non-interaction. These two models are updated iteratively until the algorithm reaches convergence. An overview of the algorithm is provided in Figure 2; the reader is referred to the Methods section for further detail.

**Fig. 2.**
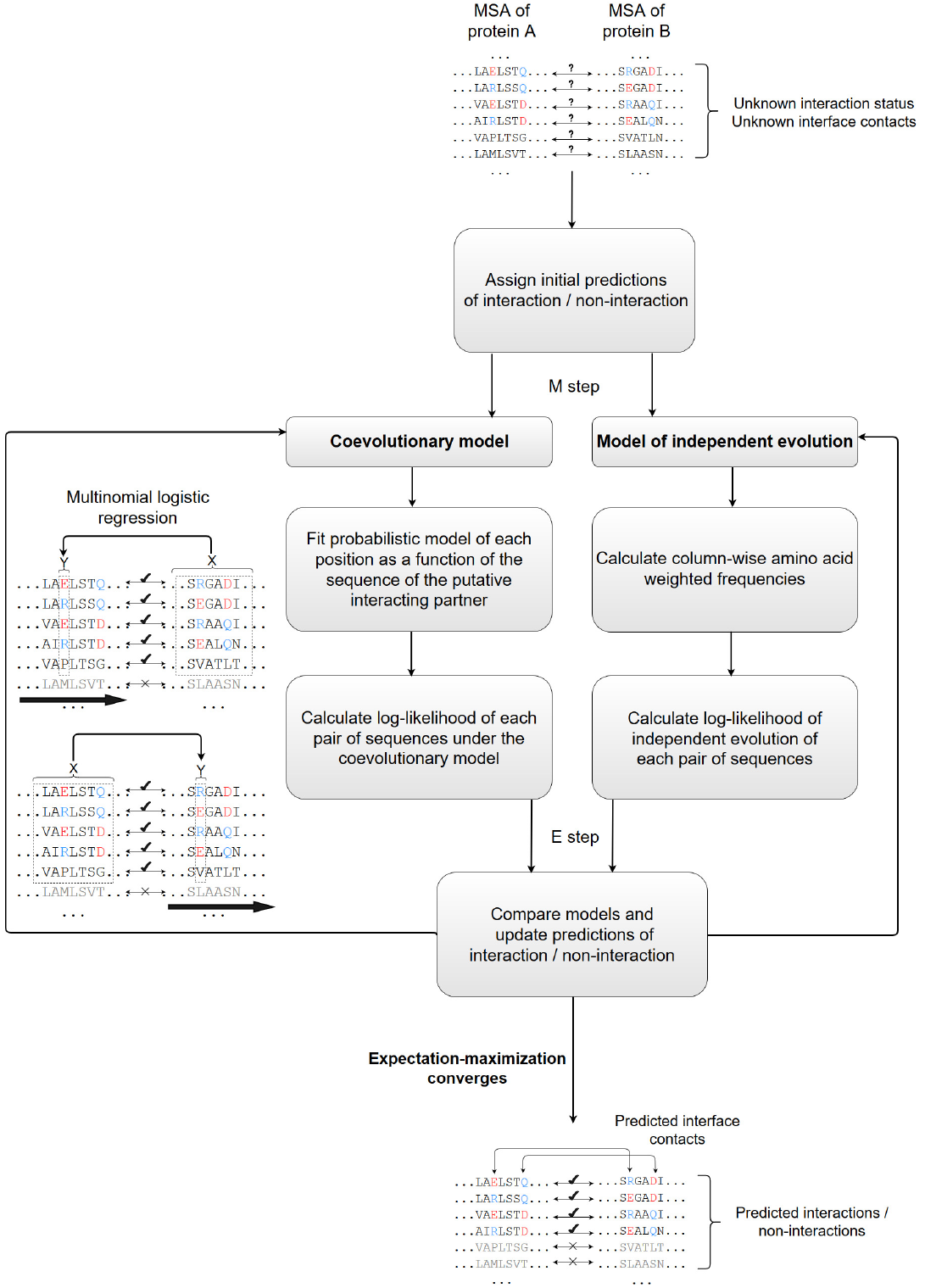
Schematic overview of the algorithm.

### F. Correlated mutations analysis accurately predicts PPIs in the PDZ-peptide dataset with no *a priori* information

We used a PDZ-peptide interaction dataset (Section D.1), which contains 82 different PDZ domains and the different peptides they bind to, to test our algorithm. As PDZ domains bind to multiple peptides, and some peptides bind to multiple PDZ domains, our alignments describe many-to-many interactions. We have created three different datasets with different levels of noise (25%, 50% and 75% of interacting proteins). All of these contain 3066 interacting sequence pairs, plus an added number of sequence pairs from our generated non-interacting set. As explained in the Methods section, there is a random element in the generation of the non-interacting set. To account for this variability, we have tested the method using datasets generated with 10 different random seeds.

Table 1 shows PPI predictive performance with different amounts of noise. A sequence pair is predicted to interact if its associated *z* value (i.e. its probability of interaction) is greater than 0.5. The case in which only 25% of the sequence pairs interact shows that very large amounts of noise can have a negative impact. Even though the algorithm still performs better than random (mean Matthews correlation coefficient of 0.17 ± 0.007; 0 would indicate no better than random performance), the predictions are rather poor compared to the other cases. In these other cases, the algorithm accurately discriminates between interacting and non-interacting sequence pairs without the need to supply knowledge about interactions, merely from covariance between MSA positions (MCCs of 0.80 ± 0.004 and 0.78 ± 0.005 for the 50% and 75% cases, respectively).

**Table 1.** PPI predictions in the PDZ-peptide dataset under different levels of noise, using 10 different random seeds. Results are presented as the mean over all random seeds and its standard error.

| % of interacting proteins | True positive rate | True negative rate | Matthews Correlation Coefficient |
| --- | --- | --- | --- |
| **25%** | $66.5 \pm 0.6\%$ | $52.2 \pm 0.7\%$ | $0.17 \pm 0.007$ |
| **50%** | $88.9 \pm 0.2\%$ | $91.1 \pm 0.4\%$ | $0.80 \pm 0.004$ |
| **75%** | $91.2 \pm 0.3\%$ | $94.5 \pm 0.3\%$ | $0.78 \pm 0.005$ |

### G. Changes in binding sites can lead to mispredictions

Intriguingly, certain domains in the interacting set are systematically misclassified as non-interacting (i.e., the median final value of *z* across all their appearances was below 0.5). These account for 5.5% of the interacting set. This consistency suggests that their sequence has some property that distinguishes them from others in the interacting set, although we found that they do not belong to a particular species or interaction specificity class. To ascertain whether the coevo-lutionary model has problems modeling certain positions, we performed principal component analysis (PCA) of the matrix of probabilities assigned by the model to the MSAs. This matrix contains the probability assigned to each individual residue in the alignments. The PCA showed some degree of separation between systematically misclassified domains and well classified domains. Most importantly, the top 3 variables with the highest absolute loading in the first principal component correspond to contact positions in the PDZ domain. These are, in our alignments, position 54, 42, and 67, by order of importance in the first principal component. Clearly, the probabilities assigned by the coevolutionary model are much lower in systematically misclassified domains than in well classified ones (Figure 3a), which indicates the coevolutionary model has problems in these three positions.

**Fig. 3.**
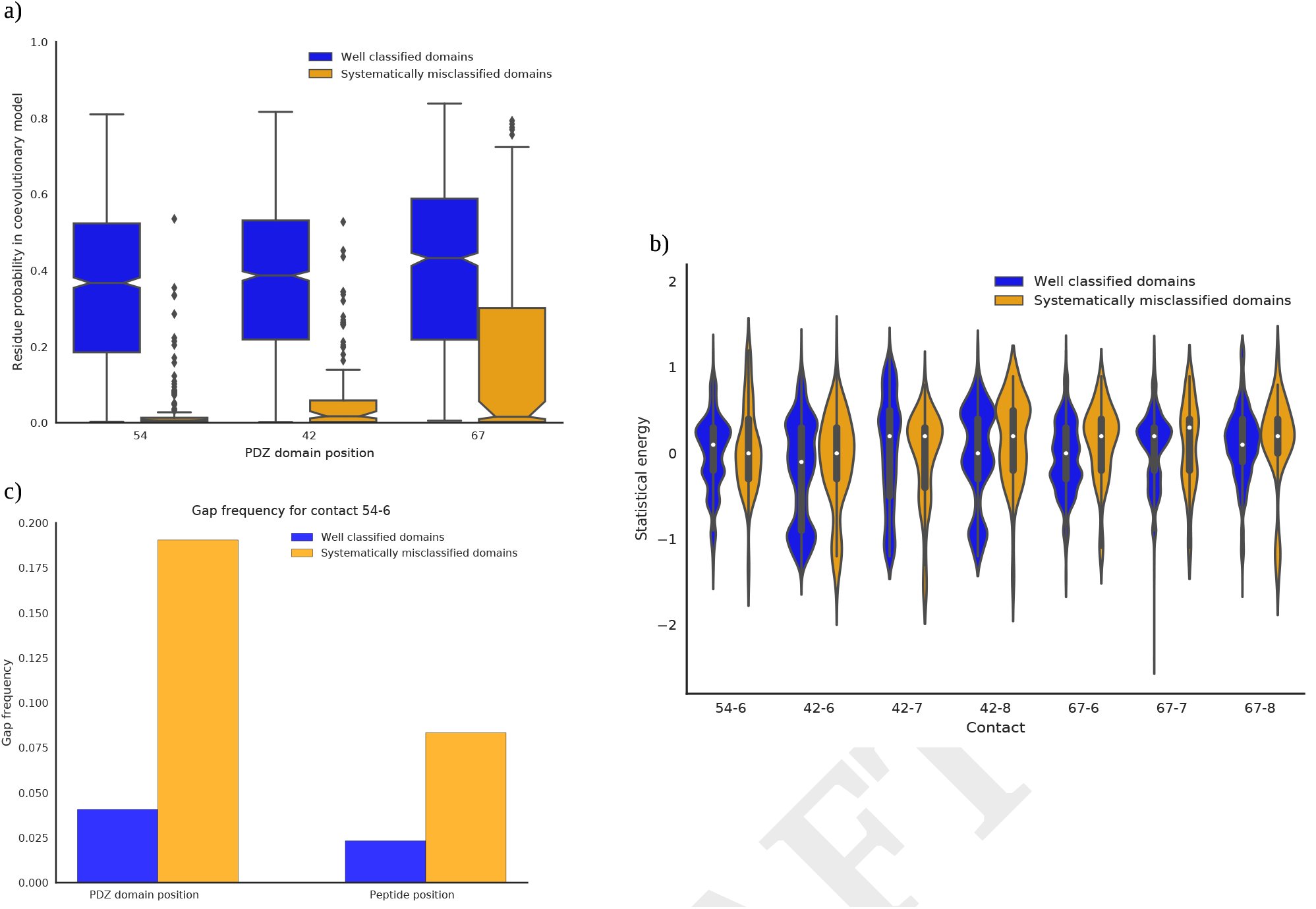
Differences between well predicted and systematically misclassified domains. (a) Boxplot showing the distribution of probabilities assigned by the revolutionary model at positions of the PDZ domain highlighted by PCA. Notches indicate the width of a 95% confidence interval of the median. (b) Violin plot illustrating distributions of statistical energies at contacts made by these positions. Boxplots are plotted within the violins. (c) Gap frequencies for the contact between domain position 54 and peptide position 6.

A general issue could be a shift in the amino acids present at the positions highlighted by PCA or in positions in the peptide that are in contact with these PDZ residues. To evaluate this possibility, we used a statistical potential (28). This potential is a function that assigns an energy to pairs of residues based on observed and expected frequencies of pairs of residues making contact in known protein structures. More negative energies between residues indicate more frequently observed, and therefore more favourable, interactions. Changes in the distribution of energies assigned by the potential would imply a change in the distribution of pairs of residues.

Figure 3b shows the distribution of these statistical energies in the different contacts made by the three PDZ residues found to behave differently according to the PCA. We find that in domain positions 42 and 67, for the consistently mis-classified sequences, there is a shift towards residue pairs with less favourable energies, especially in position 42. Position 67 still shows a higher median at all three contacts, but the effect is more moderate, as would be expected by the somewhat better probabilities assigned by the coevolutionary model (Figure 3a). However, we do not observe such a difference for position 54.

This analysis cannot take into account the presence of gaps, which could also be a problem. We observe that systematically misclassified domains have a much higher proportion of gaps at position 54 and the peptide position it interacts with (Figure 3c). Other positions did not show such an enrichment.

These factors (differences in residue pair distribution and high gap frequency in subsets of the dataset) can explain the decrease in coevolutionary model log-likelihood that leads to misclassification of this small subset of sequence pairs. Interestingly, both factors would also lead to less favourable physical interactions. It is plausible that some of these mis-classified domains bind peptides using an alternative binding mode. With our current algorithm, interactions involving such alternative binding would be difficult to detect.

### H. Accurate PPI prediction improves contact prediction ranking in noisy alignments

After we obtain PPI predictions for each sequence pair, we select those with a probability of interaction greater than 0.5. These are used to predict intermolecular contacts with plmDCA. As a baseline, we also use the alignments that contain both interacting and non-interacting proteins to predict intermolecular contacts. Also, as an ideal case, we use alignments that contain only interacting pairs. It must be noted that sequence variation, frequently measured in coevolutionary studies as the number of effective sequences, is an important factor for correct identification of contacts (see Monastyrskyy et al. 29 and Schaarschmidt et al. 30 for more details). The rather low number of effective sequences in this dataset (*N_e f f_* =76 in the set of 3066 interacting pairs, using a threshold of 80% identity) makes it somewhat difficult to identify contacts.

Figure 4 shows contact detection precision for the different datasets with increasing amounts of noise. We rank the list of predicted contacts according to their assigned coupling strength and determine the precision of the method at each point of the ranking. As previously explained, the algorithm did not manage to predict PPIs accurately in the case with only 25% interacting proteins (Figure 4a), which reflects in the poor contact prediction results. However, there is a clear improvement in the other two cases (Figure 4b, 50% interacting proteins, and Figure 4c, 75% interacting proteins), where true contacts appear earlier in the ranking. The contact prediction results obtained in noisy alignments after PPI prediction are indeed closer to those obtained by using only interacting pairs, with the last case (75% interacting proteins) following the ideal results very closely. In contrast, the naive approach of applying contact prediction to unfiltered MSAs (containing both interacting and non-interacting sequence pairs) clearly ranks the contacts worse, even in the case with 75% interacting proteins. Furthermore, the differences observed between the case with 25% interacting proteins and the others show that accurate PPI prediction is a necessary prerequisite to be able to improve contact prediction in alignments containing non-interacting sequences.

**Fig. 4.**
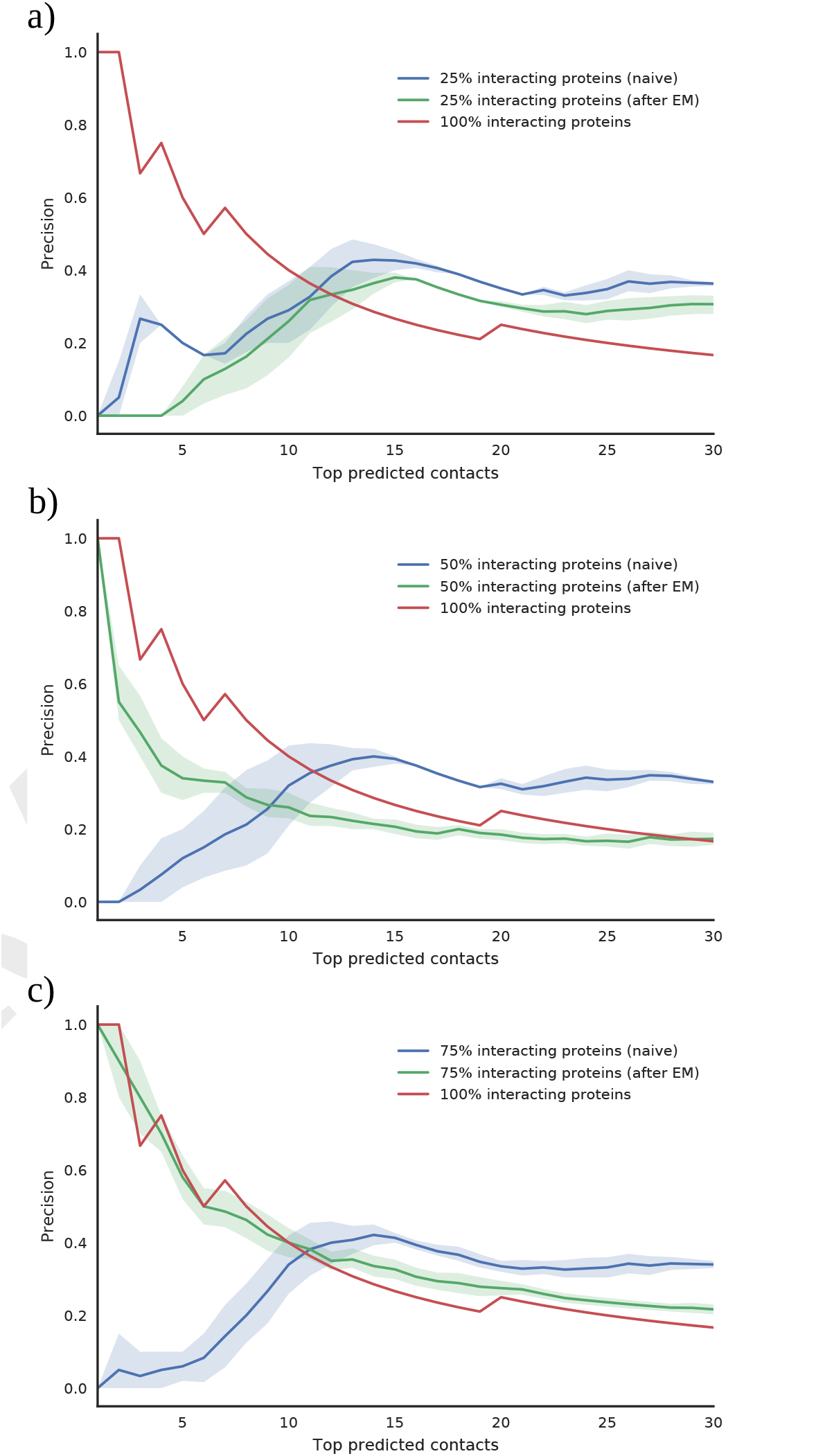
Contact prediction performance on PDZ-peptide dataset, with (a) 25% interacting proteins, (b) 50% interacting proteins, and (c) 75% interacting proteins. Naive predictions are those performed on unfiltered alignments; after EM predictions are done on alignments filtered using our method; predictions with 100% interacting proteins represent the ideal case with no non-interacting proteins. Bands indicate 90% confidence intervals of the median precision over 10 different random seeds.

### I. PPI predictions are robust to the choice of prior interacting fraction

As previously explained, the values of *z* depend on a prior fraction of interacting proteins *f_int_*. Although this parameter might be set using existing biological knowledge, we could have to set this value without any prior knowledge. In order to determine the influence that this parameter might have on the results and how it could be set without prior knowledge, we turn to our synthetic data, since we understand its underlying model and because it is free from previously mentioned confounding factors (e.g. possibly interacting negative cases). We have used synthetic data with different fractions of interacting proteins (75%, 50% and 25%), and ran the algorithm on them over a range of values for *f_int_*.

Model performance, as measured by the Matthews correlation coefficient, remains stable and close to its maximum of 1 over a wide range of settings for *f_int_*, and only deviates when the value is quite far from the true value (Figure 5a) for all three datasets. Note that here, in contrast to results obtained on the PDZ-peptide dataset, we obtain good predictive performance when we only have 25% of interacting proteins. One possible reason is the larger amount of sequence variation in this dataset (*N_e f f_*=550 at a threshold of 80% identity, using only the interacting set).

**Fig. 5.**
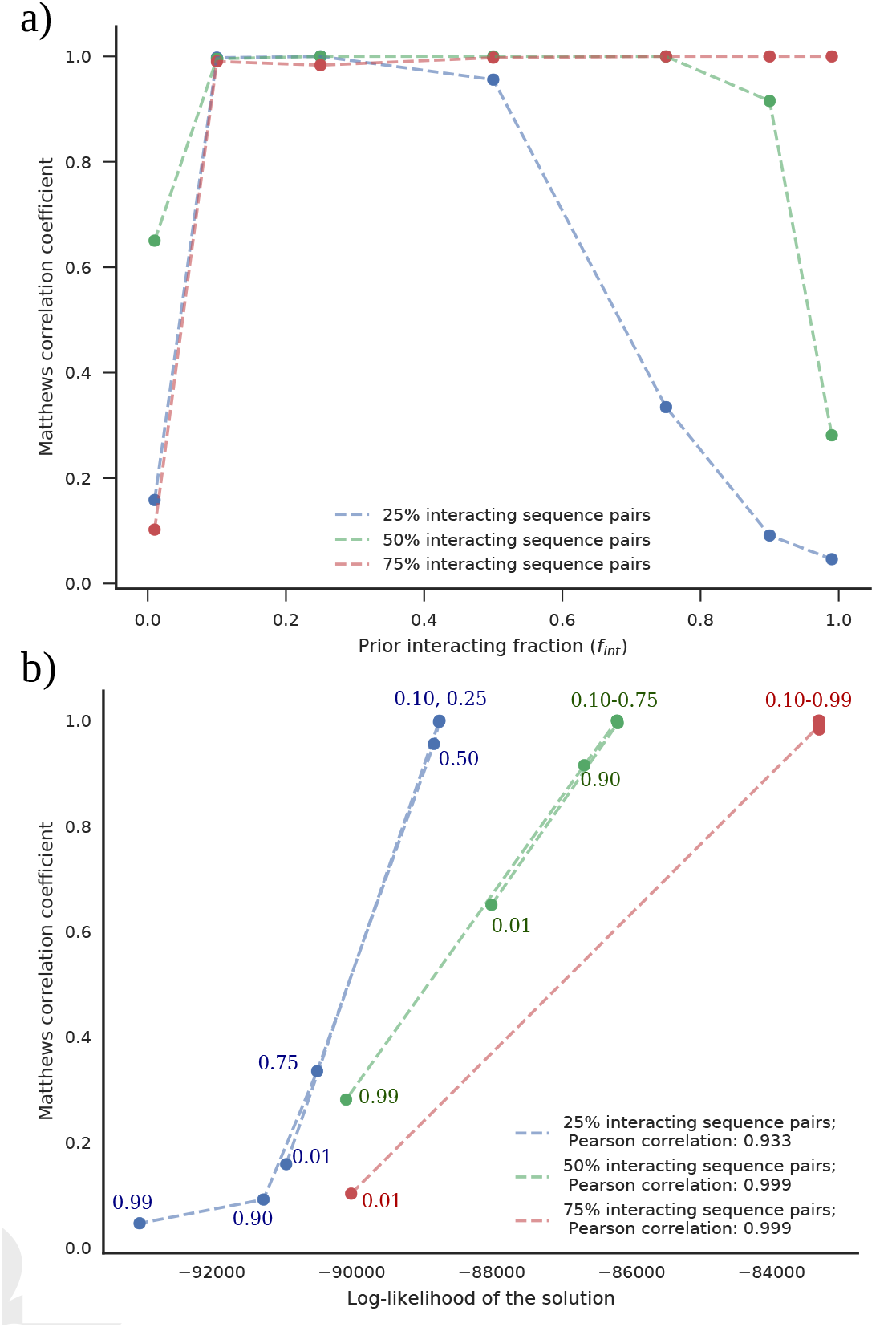
Results on synthetic MSAs. (a) Model performance remains stable over a wide range of settings for *f_int_*. (b) Model performance peaks together with the log-likelihood of the data in all three datasets. The value of *f_int_* is indicated next to the points. The asterisk and the triangle indicate all used values of *f_int_* from 0.10 to 0.75 (50% interacting proteins) and from 0.10 to 0.99 (75% interacting proteins), respectively.

Furthermore, model performance correlates extremely well with the log-likelihood of the solution reached by the algorithm (which is a function of the model parameters, and can be computed without prior knowledge of protein-protein interaction) (Figure 5b). Thus, in the absence of prior knowledge, the value of *f_int_* can be set in a principled way by testing several values and choosing the one that yields the maximum log-likelihood.

## Conclusions

We have designed Ouroboros, a combination of coevolutionary analysis and EM to approach the problem of predicting intermolecular contacts using MSAs containing both interacting and non-interacting proteins. We have shown that the method is able to precisely predict the interaction status between two given proteins. This, in turn, allows us to reach our goal of improving contact prediction performance in noisy MSAs.

Although here we have focused on the use of sequence data, other sources of information could be incorporated to guide the algorithm. For instance, if the interaction status between certain sequence pairs in the alignments is known, this could be used as a constraint for EM (see, for example, Ganchev et al. 31). This could make it reach convergence faster and improve PPI prediction performance (and, by extension, contact prediction performance). This would be particularly relevant for cases that contain a large number of non-interacting sequence pairs or where there is low sequence diversity. Likewise, knowledge on interaction sites (based, for example, on mutagenesis data) may also prove useful.

We have also found that high gap frequency and substitutions at contact positions in subsets of the MSAs can be a confounding factor and lead to PPI misclassification. It is possible that pairs of proteins where this occurs have developed a different binding mode, with other positions compensating the loss of contacts. In a similar fashion, it has been highlighted that subfamily-specific signals can be a confounding factor in coevolutionary analysis of homomeric interactions (32). The concepts and methodology detailed in this paper could be further developed to accomplish an additional objective related to these confounding factors: distinguishing between subgroups of proteins in either type of interaction, thereby revealing these specific signals without prior knowledge of the subgroups.

In addition, although we have focused on the task of contact prediction, coevolution-based methods for PPI prediction offer an interesting alternative to methods that rely on sequence similarity. This is especially important for the study of protein families with similar sequences but highly divergent interaction patterns.

In summary, the developed algorithm can successfully improve contact prediction in MSAs with non-interacting proteins without *a priori* knowledge of interactions. Furthermore, it allows for many-to-many interactions, and thus opens the way for the study of interactions that were previously out of reach.

